# Human organoid model of PCH2a recapitulates brain region-specific pathology

**DOI:** 10.1101/2022.10.13.512020

**Authors:** Theresa Kagermeier, Stefan Hauser, Kseniia Sarieva, Lucia Laugwitz, Samuel Groeschel, Wibke Janzarik, Zeynep Yentür, Katharina Becker, Ludger Schöls, Ingeborg Krägeloh-Mann, Simone Mayer

**Author notes:** Corresponding author: Dr. Simone Mayer, Hertie Institute for Clinical Brain Research, Otfried-Müller Str. 25, 72076 Tübingen, Germany.

## Abstract

Pontocerebellar hypoplasia type 2 a (PCH2a) is a rare, autosomal recessive pediatric disorder with limited treatment options. Its anatomical hallmark is the hypoplasia of the cerebellum and pons accompanied by progressive microcephaly. PCH2a results from a homozygous founder variant in *TSEN54*, which encodes a tRNA splicing endonuclease (TSEN) complex subunit. Despite the ubiquitous expression of the TSEN complex, the tissue-specific pathological mechanism of PCH2a remains unknown due to a lack of model system. In this study, we developed human models of PCH2a using brain region-specific organoids. We therefore obtained skin biopsies from three affected males with genetically confirmed PCH2a and derived induced pluripotent stem cells (iPSCs). Proliferation and cell death rates were not altered in PCH2a iPSCs. We subsequently differentiated cerebellar and neocortical organoids from control and PCH2a iPSCs. Mirroring clinical neuroimaging findings, PCH2a cerebellar organoids were reduced in size compared to controls starting early in differentiation. We observed milder growth deficits in neocortical PCH2a organoids. While PCH2a cerebellar organoids did not upregulate apoptosis, their stem cell zones showed altered proliferation kinetics, with increased proliferation at day 30 and reduced proliferation at day 50 compared to controls. In summary, we have generated a human model of PCH2a, which provides the foundation for deciphering brain region-specific disease mechanisms.

## Introduction

Pontocerebellar hypoplasias (PCH) comprise a heterogeneous group of neurogenetic disorders characterized by a severe neurodevelopmental impairment and hypoplasia of the cerebellum and pons (van Dijk et al 2018). Besides the primary pontocerebellar hypoplasia, affected individuals develop progressive microcephaly (Barth et al 2007, Namavar et al 2011). PCH2 is the most common form of PCH but is still ultra-rare (1:100,000 births) (Budde et al 2008, Sanchez-Albisua et al 2014). Clinically, PCH2 is characterized by profound neurodevelopmental delay, causing a significant burden to affected individuals and their families (Ammann-Schnell et al 2021, Sanchez-Albisua et al 2014). To date, no curative treatment is available, and disease management is focused on alleviating symptoms.

90% of PCH2 cases are caused by a homozygous founder variant in the *TSEN54* gene (OMIM *608755) (NM_207346.3:c.919G>T, p.(Ala307Ser)) and are referred to as PCH2a (OMIM #277470) (Budde et al 2008, van Dijk et al 2018). *TSEN54* encodes a subunit of the tRNA splicing endonuclease (TSEN) complex, which is involved in excising introns from a subset of pre-tRNAs (Chan & Lowe 2009, Trotta et al 2006). The TSEN54 protein region that contains Ala307Ser variant is not visualized by cryo-EM, thus suggesting that it is unstructured (Hayne et al 2023, Sekulovski et al 2023). TSEN54 is thought to be a structural subunit of TSEN complex (Hayne et al 2023, Sekulovski et al 2023). While bi-allelic pathogenic variants in *TSEN54* do not affect the endonuclease activity of the TSEN complex, in fibroblasts of affected individuals, they result in a thermal destabilization of the complex and altered tRNA pools (Sekulovski et al 2021). The TSEN54 protein is expressed throughout the body at varying levels (Human Protein Atlas) (Uhlen et al 2015). At the mRNA level, *TSEN54* is widely expressed in the developing human brain (Human Brain Transcriptome) (Kang et al 2011) starting in the first trimester of gestation (Kasher et al 2011). In the second trimester of gestation, *TSEN54* is expressed highly in the developing cerebellum, pons, and olivary nuclei (Budde et al 2008). Its expression does not appear to be cell type-specific in the developing neocortex and cerebellum at the level of mRNA (Aldinger et al 2021, Nowakowski et al 2017). Collectively, from the TSEN54 expression pattern alone, it is unclear why specifically the cerebellum and pons are affected in PCH2a (Kasher et al 2011). Interestingly, additional variants in *TSEN54* (PCH4,5), as well as variants in other subunits of the TSEN complex(PCH2b,c,f),, and variants in *CLP1* (PCH10), which encodes a protein that interacts with the TSEN complex, result in clinical phenotypes related to PCH2a (Schaffer et al 2019). Therefore, it has been hypothesized that specific brain areas are especially vulnerable to TSEN malfunction due to a specific requirement of this complex during early postnatal development (Budde et al 2008).

Histopathological analysis to date has revealed a reduced complexity of cerebellar foliation, patches of missing cerebellar cortex lacking Purkinje cells (PC) and severely reduced and misplaced granule cells (GC), the major neuronal cell types of cerebellar cortex (Rudnik-Schöneborn et al 2014). Efforts to recapitulate the cellular mechanism of pathophysiology in animal models have been inconclusive so far. While the TSEN complex is conserved from archaea, *TSEN54* has undergone evolutionary changes in the primate lineage (Lee et al 2016). Notably, the amino acid sequence around *TSEN54*:c.919G>T, p.(A307S) is not conserved between species (Budde et al 2008). Of the frequently used model organisms, only mouse and chicken share the amino acid residue (Budde et al 2008). A recently developed fly model of PCH shows defects in brain development, death at larval stages, and apoptosis upon loss of function of the *TSEN54* ortholog, but the relevance to the brain region-specific clinical phenotype is unclear (Schmidt et al 2022). In zebrafish, loss of *tsen54* function leads to cell death in the brain during development (Kasher et al 2011). A complete loss of *Tsen54* in mice results in early embryonic lethality (Ermakova et al 2018). Moreover, bi-allelic missense variant of *TSEN54* in dogs leads to a neurological disorder characterized by leukodystrophy, a disease with a strikingly different pathophysiology (Stork et al 2019). These findings imply a potential species-specific effect of TSEN54 malfunction on brain development. Additionally, the brain has changed tremendously in evolution, especially in the primate lineage (Herculano-Houzel 2012). Compared to mouse, the human neocortex has expanded dramatically (1000x in the number of neurons and surface area), has a protracted development (neurogenesis is 20x longer than in mouse) and displays an unprecedented cellular heterogeneity (Geschwind & Rakic 2013, Miller et al 2019). Similarly, the human cerebellum has a 750-fold greater surface area than the mouse, a protracted development occurring over 2-3 years compared to 30-35 days, and additional transient stem cell zones that cannot be found in mouse (Haldipur et al 2022). Taken together, the lack of appropriate models of PCH2a has to date precluded the elucidation of its cellular and molecular pathology.

As an alternative to animal models, human brain organoid models have recently been increasingly used to model neurodevelopmental and early-onset neurodegenerative disorders as well as investigating environmental impacts on brain development (Eichmüller & Knoblich 2022, Sarieva & Mayer 2021). In order to study brain region-specific biology, it is possible to guide the differentiation of organoids to a specific brain region by adding specific morphogens, (Kadoshima et al 2013).. Neocortical organoids, for instance, recapitulate cell type composition of the developing human neocortex with SOX2+ ventricular radial glia cells organized into rosettes reminiscent of ventricular zone (VZ) and shed other neural progenitor cells (NPC) and neurons to putative subventricular zone (SVZ) and cortical plate (CP), respectively (Pasca et al 2015). Within the CP-like regions, a distinct lamination pattern emerges, featuring CTIP2+ deep-layer (dExN) and SATB2+ upper-layer (uExN) excitatory neurons (Pasca et al 2015).

Similarly, in cerebellar organoid differentiation protocols (Muguruma et al 2015, Silva et al 2020), the stem cells of two cerebellar NPC zones are established: KIRREL2+/PFT1A+ VZ and BARHL+/ATOH1+ rhombic lip (RL) (Leto et al 2016, Mizuhara et al 2010). These NPCs generate neurons of two major cerebellar neuronal identities: PCs and GCs. PC precursors (KIRREL2+/PFT1A+) derive from the VZ, and are identified by OLIG2/SKOR2 expression and subsequently mature into CALB+ PCs. (Leto et al 2016, Wang et al 2011). RL derived precursors (BARHL+/ATOH1+) differentiate into GCs and glutamatergic deep cerebellar nuclei (DCN) neurons, which are positive for LHX2 (Ballabio et al 2020, Kamei et al 2023, Lago et al 2023, Morales & Hatten 2006).

In this study, we leverage the recent developments in the generation of brain region-specific organoids (Zhang et al 2022) to create a human model of PCH2a. We identified three individuals that display the genetic, clinical and brain imaging features previously described for PCH2a and derived induced pluripotent stem cells (iPSCs) from donated fibroblasts (Figure 1A). We did not find differences in proliferation and cell death between PCH2a and control iPSCs. Next, we differentiated PCH2a and control iPSC towards a cerebellar and neocortical fate in organoid cultures. Growth curves of organoids recapitulated the brain region-specific pathology observed in affected individuals. Cerebellar PCH2a organoids were severely reduced in size compared to controls starting early in differentiation, while neocortical PCH2a organoids showed less severe divergence from controls only at later stages of differentiation. We did not detect differences in the induction of apoptosis in NPCs. Instead, we find an increased number and thickness of neural rosettes along with increased proliferation in these areas in early but not late differentiation stages in PCH2a cerebellar organoids compared to controls. Our study thus provides first insights into disease mechanisms, which can be analyzed in depth in future studies by using the organoid model of PCH2a established here.

**Figure 1:**
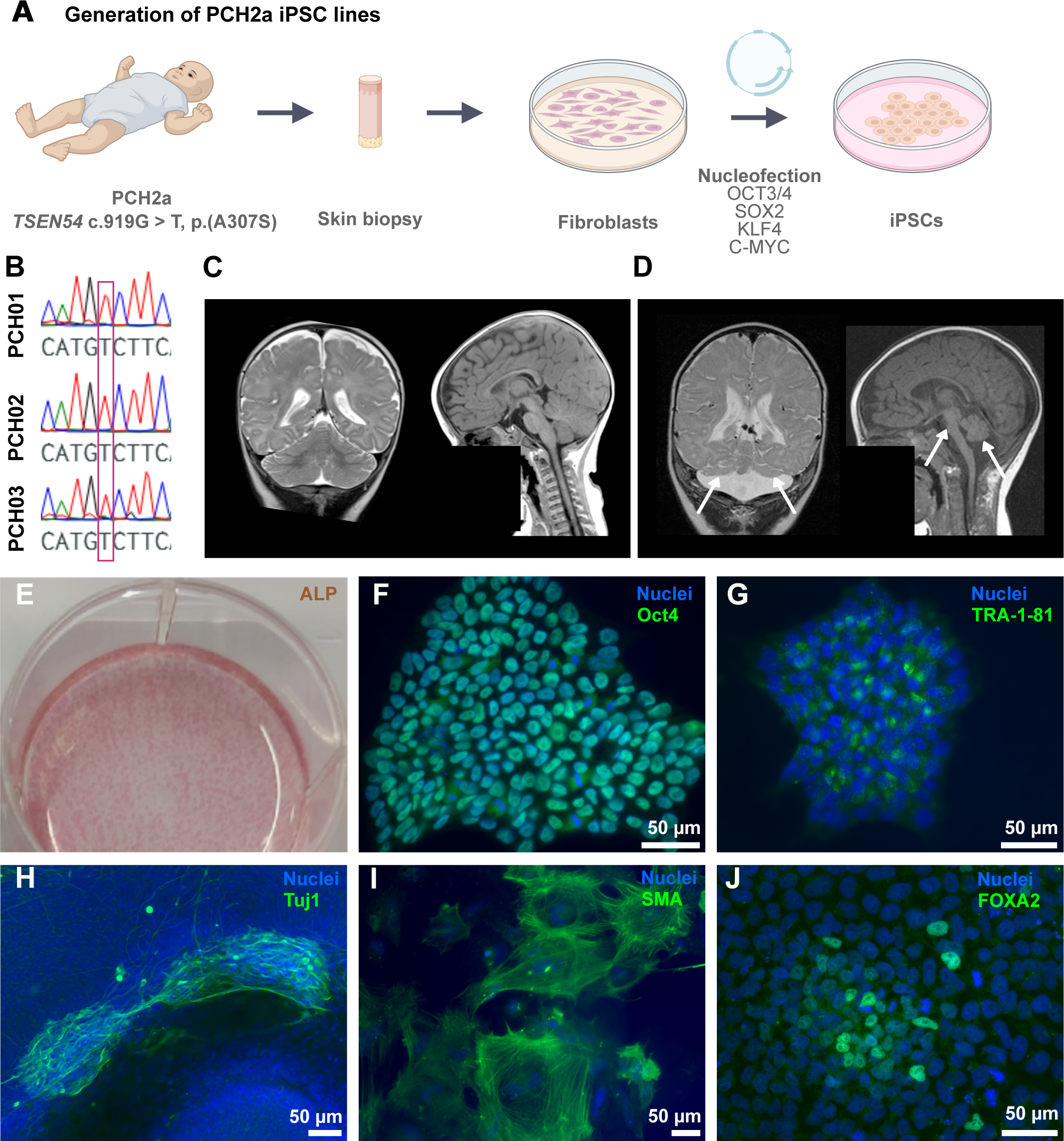
Generation of PCH2a iPSCs. (A) Experimental scheme of PCH2a iPSC generation. (B) Sanger sequencing results of generated PCH2a iPSCs verifies c.919G>T point mutation of PCH2a-derived iPSCs. (C) MRI of the brain of a normally developing infant (6 months old) and (D) an infant with PCH2a (6 months old, donor of iPSC line PCH02). T2w coronal images (C and D, left) show cerebellar hemispheres severely reduced in size (indicated by arrows) in the PCH2a child compared to the control individual. T1w sagittal images (C and D, right) illustrate the severe pontine and cerebellar hypoplasia in the PCH2a child (indicated by arrows). The facial region of the MRI scans are covered in order to protect the privacy of both individuals. (E) Alkaline phosphatase (ALP) staining of undifferentiated iPSCs. Immunocytochemical stainings of undifferentiated iPSCs for the pluripotency markers OCT4 (F) and TRA1-81 (G) and DAPI (nuclei) demonstrate the pluripotency of generated cell lines. (H-J) Immunocytochemical staining of iPSCs spontaneously differentiated into the three germ layers illustrates the differentiation potential of the generated iPSCs. Cells were stained for Tuj1 (ß-III-tubulin, ectoderm). (H), SMA (smooth muscle actin, mesoderm) (I), and FOXA2 (Forkhead Box A2, endoderm) (J), as well as DAPI (nuclei). Representative images show PCH01.

## Results

### Generation of PCH2a-derived iPSCs

It is unclear how a ubiquitously expressed gene involved in tRNA metabolism causes pathology only in specific tissues and even within the nervous system, with differential pathology in different brain regions. Based on recent successes in modeling neurogenetic disorders in organoids (Khakipoor et al 2020, Velasco et al 2020), we reasoned that a human iPSC-based brain region-specific organoid model could recapitulate pathological hallmarks of PCH2a. Previously, cerebellar organoids were used to investigate the mechanisms of pediatric cerebellar cancer, medulloblastoma (Ballabio et al 2020, Lago et al 2023). We hypothesized that generating neocortical and cerebellar organoids might model the brain region-specific pathology in PCH2a. However, to our knowledge, cerebellar organoids have not yet been used to model neurogenetic disorders involving the cerebellum.

As a first step, we identified three affected males with genetically confirmed PCH2a, harboring the c.919G>T variant in the *TSEN54* gene in the homozygous state (Figure 1B). Clinical features typical of PCH2a were evident in all probands (Sanchez-Albisua et al 2014). In summary, subjects displayed severe developmental delay with minimal cognitive and motor development, seizures of variable semiology, and a severe dystonic movement disorder. Moreover, the probands exhibited neurogenic dysphagia and gastrointestinal disturbances (Supplementary Table 1). Neuroimaging revealed severe hypoplasia of the brainstem, pons and cerebellum in contrast to less severe volume reduction of the cerebrum as seen in affected child (donor of iPSC line PCH02) (Figure 1C) in comparison to an age-matched control (Figure 1D).

We obtained skin biopsies from these three probands, extracted fibroblasts and derived iPSCs using an episomal reprogramming approach (Figure 1A) (Okita et al 2013). iPSCs were subjected to a range of quality controls. First, pluripotency was confirmed through staining for alkaline phosphatase (ALP) (Figure 1E) and immunocytochemistry for OCT4 and TRA-1-81 (Figure 1F,G). Differentiation potential was corroborated using spontaneous tri-lineage differentiation (Korneck et al 2022). iPSCs differentiated into the three primordial germ layers, namely, ectoderm (Tuj1), mesoderm (SMA), and endoderm (FOXA2) (Figure 1H-J). Control lines used in this study were generated following the same protocol and were subjected to all aforementioned quality controls (see methods section for detailed description). SNP array analysis of all six lines used in this study revealed no larger chromosomal aberrations induced by the reprogramming (Supplementary Data 4).

In order to determine even subtle differences in iPSC properties between PCH2a and control lines, we assessed and quantified the number of cells expressing pluripotency marker OCT4 (Figure 2A,B), proliferative marker Ki-67 (Figure 2C,D) and apoptotic marker cleaved caspase-3 (cCas3) (Figure 2E,F) of three consecutive passages of all PCH2a and control iPSC lines used in this study. We identified no significant differences in the number of cells expressing these markers between PCH2a and control lines. Additionally, we incubated the iPSC lines with EdU for 1 and 4 hours to assess the proliferation rate (Figure 2G,H). Quantification of EdU+ cells revealed no significant difference in EdU incorporation between PCH2a and control lines. These results indicate that iPSCs derived from affected individuals maintain characteristics of control iPSC lines and hence can be used for studying tissue-specific pathology by employing region-specific organoid differentiation protocols. In conclusion, the pathogenic variant does not seem to affect the proliferation and viability of the iPSCs.

**Figure 2:**
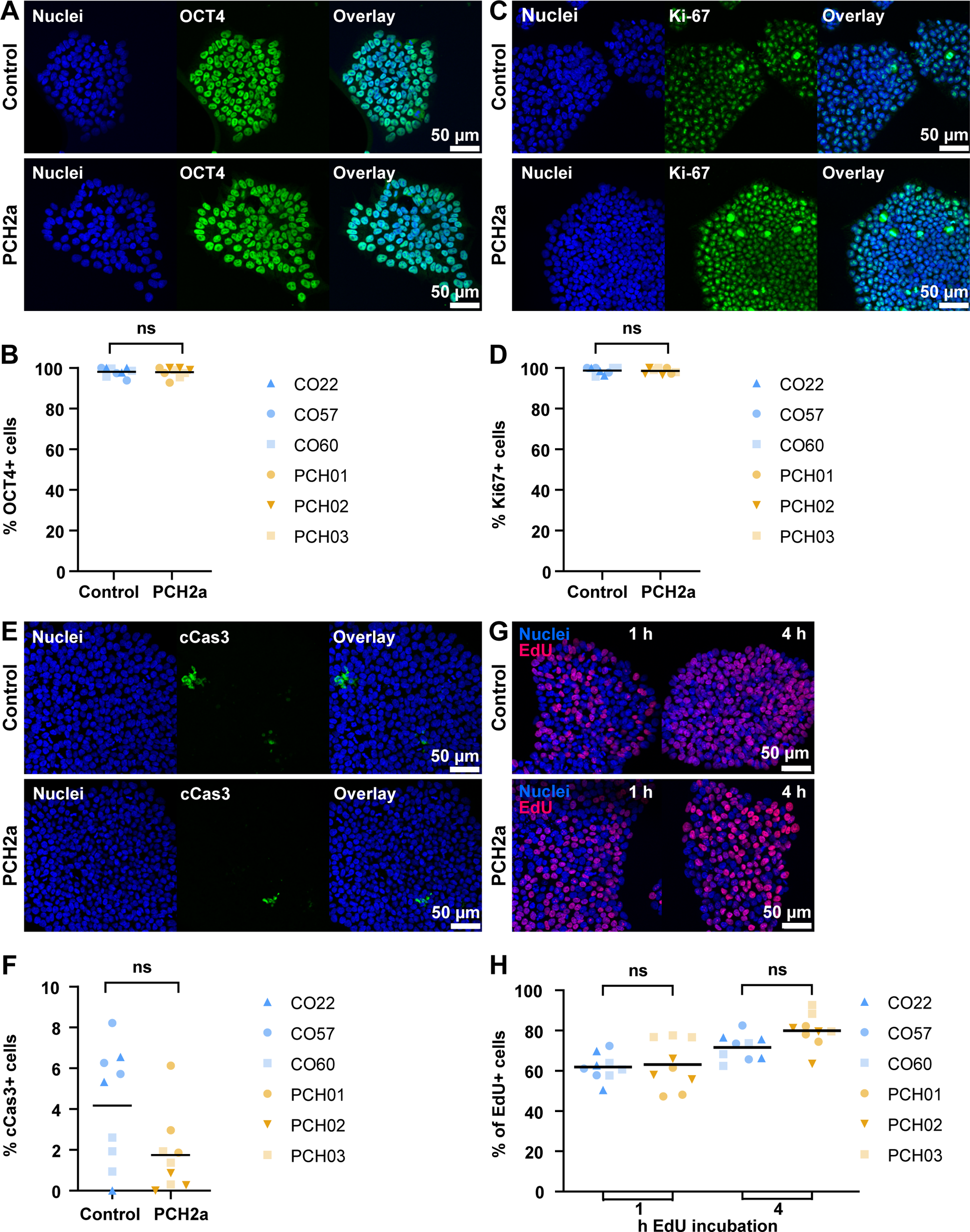
PCH2a-derived and control iPSCs do not differ in expression of pluripotency, proliferation and apoptosis markers. (A) Immunocytochemical staining of PCH2a and control iPSCs for pluripotency marker OCT4 confirms the expression in all iPSC lines (representative image CO22 P19, PCH01 P17). (B) Quantification of OCT4 positive cells, normalized to DAPI-based cell count shows no significant difference in number of OCT4+ cells between PCH2a and control iPSCs (assessment of 3 passages per cell line). (C) Immunocytochemical staining of PCH2a and control iPSCs for proliferation marker Ki-67 confirms the expression in all iPSC lines (representative image CO57 P20, PCH03 P20). (D) Quantification of Ki-67 positive cells, normalized to DAPI-based cell count shows no significant difference in number of Ki-67+ cells between PCH2a and control iPSCs (assessment of 3 passages per cell line). (E) Immunocytochemical staining of PCH2a and control iPSCs for apoptosis marker cCas3 shows low expression levels in all iPSC lines (representative image CO60 P18, PCH02 P18). (F) Quantification of cCas3 positive cells, normalized to DAPI-based cell count shows no significant difference in number of cCas3+ cells between PCH2a and control iPSCs (assessment of 3 passages per cell line). (G) Click-chemistry detection of EdU on PCH2a and control iPSCs after 1 and 4 hours of incubation with EdU visualizes cell proliferation (representative image CO60 P18, PCH02 P18). (H) Quantification of EdU positive cells, normalized to DAPI-based cell count after 1 and 4 hours of EdU incubation shows no significant difference in number of EdU positive cells between PCH2a and control iPSCs (assessment of 3 passages per cell line) (ns, p>0.05 unpaired t-test with Welch’s correction assuming unequal SDs).

### Human brain organoids have brain region-specific growth deficits

To model brain region-specific pathology, we then differentiated three control and the three PCH2a iPSC lines towards a cerebellar and neocortical fate in 3D using established protocols (Pasca et al 2015, Silva et al 2020) (Figure 3A).

**Figure 3:**
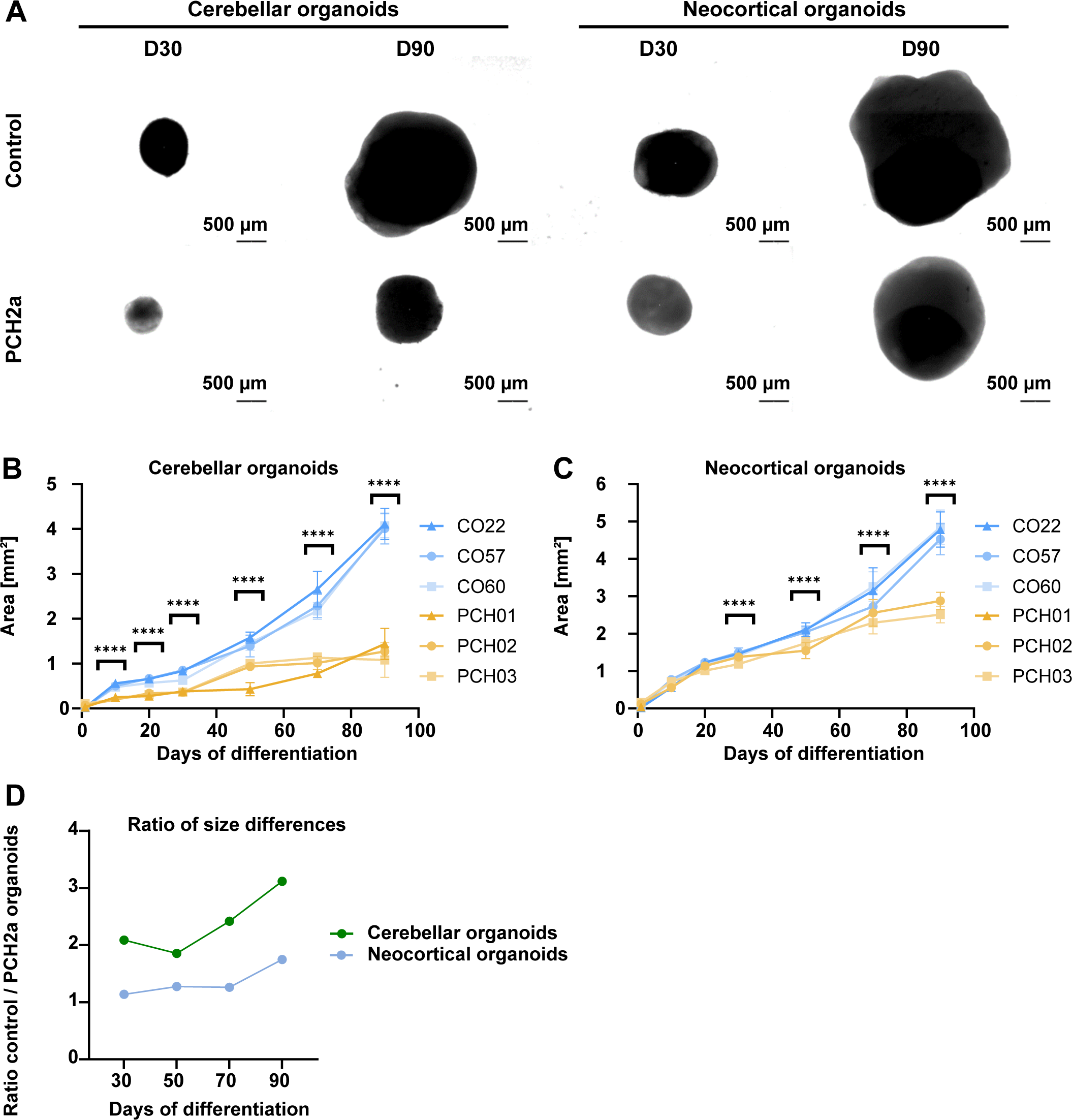
PCH2a organoids are significantly smaller than control organoids. (A) Representative bright field images of cerebellar and neocortical organoids in culture at D30 and D90 of differentiation illustrate the differences in size. (B, C) Growth curves of PCH2a and control cerebellar (B) and neocortical (C) organoids, differentiated from 3 different cell lines per condition (derived from 3 different individuals), show the area of the organoids in the images pictured in A during the culture period of 90 days. Cerebellar organoids (B) differ significantly in size from D10 of differentiation (p< 0.05 unpaired t-test with Welch’s correction assuming unequal SDs), with discrepancies increasing over time. Neocortical organoids (C) show significant differences from D30 of differentiation (p< 0.05 unpaired t-test with Welch’s correction assuming unequal SDs). n>8 organoids per cell line, timepoint and differentiation. Points represent the mean, error bars represent SEM. Note that PCH01 neocortical differentiation is absent due to a contamination. (D) Ratio between mean sizes of control / PCH2a organoids calculated from data presented in (B) and (C). Ratios increase over time and are higher within cerebellar differentiation.

We first determined if cerebellar hypoplasia and progressive microcephaly found in affected individuals (Figure 1C,D) were recapitulated in our *in vitro* model (Figure 3B,C). We, therefore, measured the sizes of both cerebellar and neocortical organoids in brightfield images at day (D) 10, D20, D30, D50, D70, and D90. During neocortical differentiation one line (PCH2a) was contaminated and the differentiation was terminated at D20. Interestingly, for both brain region-specific organoid differentiations, we found significant differences in size between PCH2a and control organoids (Figure 3A-C, Supplementary Figure 1A-F). On average, cerebellar PCH2a organoids were smaller than controls starting from D10 of differentiation (Figure 3B), while neocortical PCH2a organoids showed differences from D30 onwards (Figure 3C). Statistical assessment (3-way ANOVA) revealed no significant difference for the sizes within PCH2a or control groups at any time point justifying the comparison of mean sizes between groups. Linear regression models were significantly different between control and PCH2a organoids of both regional identities (Supplementary Figure 1G,H, supplementary table 2). At D50, the ratio between the mean sizes of control / PCH2a cerebellar organoids was 1.85, increasing to 3.12 at D90 (Figure 3D). In neocortical organoids, the ratio between the mean sizes of control and PCH2a organoids was 1.13 at D50 and 1.74 at D90 of differentiation (Figure 3D). The brain region-specific organoid growth curves resemble brain morphometry in PCH2a-affected individuals with early detection of cerebellar hypoplasia in the first months of life (Sanchez-Albisua et al 2014) and the later progressive microcephaly (Ekert et al 2016), albeit at a different time scale.

### PCH2a-derived iPSCs differentiate towards cerebellar and neocortical fate

To start assessing the cellular underpinnings of the brain region-specific growth deficits (Figure 3), we analyzed the presence of different progenitor and neuronal cell types in cerebellar and neocortical organoids at different time points in PCH2a and control organoids. We found robust neural differentiation in both PCH2a and control cerebellar organoids based on immunohistochemical analysis for the markers SOX2 (NPCs) and Tuj1 (immature neurons) at D30 (Figure 4A,B). Moreover, cerebellar organoids demonstrated presence of both RL- and VZ-derived cerebellar NPC populations. Immunohistochemistry against BARHL1 (Figure 4A,B, Supplementary Figure 2A,B) and ATOH1 (Supplementary Figure 3A-D) confirmed the presence of RL-derived glutamatergic cerebellar precursor cells. Cerebellar VZ derivates such as GABAergic cerebellar precursors are characterized by KIRREL2 (Figure 4C,D), OLIG2 (Supplementary Figure 2E,F) and SKOR2 (Supplementary Figure 3A-D) in immunohistochemistry at D30. At later stages of differentiation (D90), we found Calbindin (CALB) and MAP2-positive cells, indicating the presence of PCs in cerebellar organoids (Figure 4E,F). Similarly, in neocortical organoids, immunohistochemical analysis for CTIP2, SATB2, SOX2, and Tuj1 collectively revealed neocortical differentiation and generation of layer-specific excitatory neurons (Supplementary Figure 3A-F). We found expression of the NPC marker SOX2 in neural rosettes surrounded by CTIP2-positive dExNs at D50 in control (Supplementary Figure 3A) and PCH2a (Supplementary Figure 3B) neocortical organoids. In addition, expression of the uExN marker SATB2 was found at D70 and D90 (Supplementary Figure 3C-F). Taken together, the acquisition of brain region-specific fate and neuronal maturation was evident in both cerebellar and neocortical control and PCH2a organoids, suggesting they can be used to analyze molecular and cellular changes induced by the disease-causing variant.

**Figure 4:**
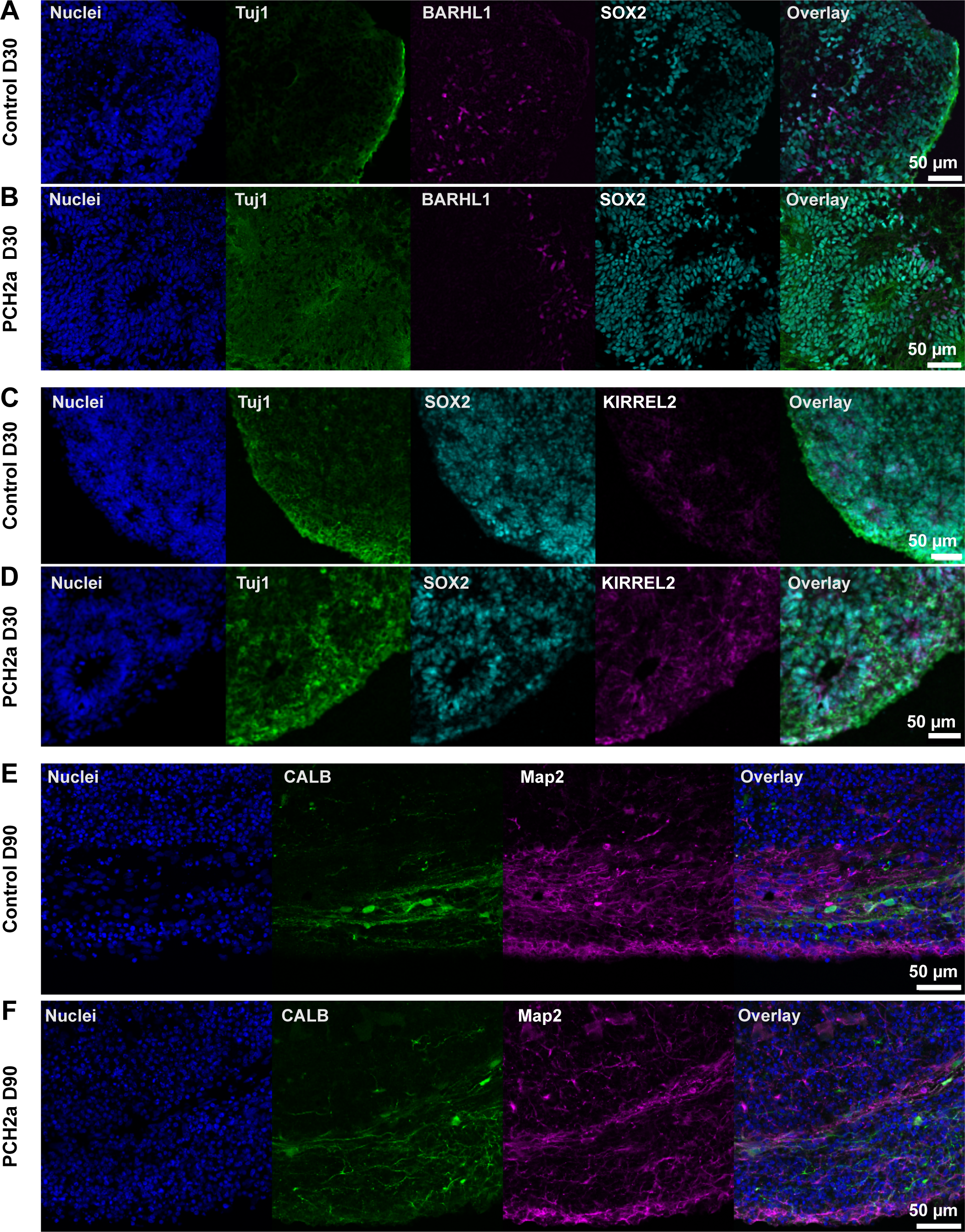
PCH2a and control cerebellar organoids show differentiation into cerebellar lineage. Immunohistochemistry of control and PCH2a cerebellar organoid sections at D30 and D90 show differentiation into cerebellar lineage. (A, B) Expression of early neuronal marker Tuj1 (green), neural precursor marker SOX2 (cyan), and glutamatergic precursor marker BARHL1 (magenta) in D30 control (A) and PCH2a (B) cerebellar organoids (representative images CO22 P19, PCH02 P17). (C, D) GABA-ergic precursor marker KIRREL2 (magenta) is expressed in D30 cerebellar control (C) and PCH2a (D) organoids together with Tuj1 (green) and precursor marker SOX2 (cyan) (representative images CO22 P19, PCH01 P19). (E, F) Control (I) and PCH2a (J) cerebellar organoids show Calbindin (CALB, cyan) in postmitotic Purkinje cells and neuronal marker MAP2 (magenta) expression (representative images CO57 P18, PCH03 P19).

### Expression of apoptotic marker cleaved Caspase-3 is not altered in PCH2a cerebellar and neocortical organoids

Previous histopathological assessment of a PCH2a affected individual suggested a degenerative nature of the disease (Barth et al 2007, Rudnik-Schöneborn et al 2014). Additionally, studies on zebrafish and fruit fly TSEN54 ortholog loss of function models indicated that hypoplasia resulted from cell death (Kasher et al 2011, Schmidt et al 2022). We therefore investigated whether elevated levels of apoptosis could explain the reduced size of PCH2a cerebellar and neocortical organoids. However, immunohistochemistry for the apoptotic marker cCas3 did not reveal differences between control and PCH2a cerebellar (Figure 5A) and neocortical (Figure 5B) organoids. Quantitative analysis of the cCas3-positive area over the DAPI area of individual regions of interest (SOX2-rich regions containing SOX2+ rosettes and surrounding cells) did not show a significant difference between PCH2a and control organoids at D30 and D50 in both cerebellar (Figure 5C) and neocortical (Figure 5D) organoids. Further quantification of the cCas3+ area within the SOX2+ area only did not show an elevated apoptosis rate in the SOX2+ NPC population (Figure 5E,F). Taken together, we observe a significant size difference in PCH2a brain organoids (Figure 3) in the absence of obvious changes in apoptosis at D30 and D50 of differentiation in cerebellar and neocortical organoids.

**Figure 5:**
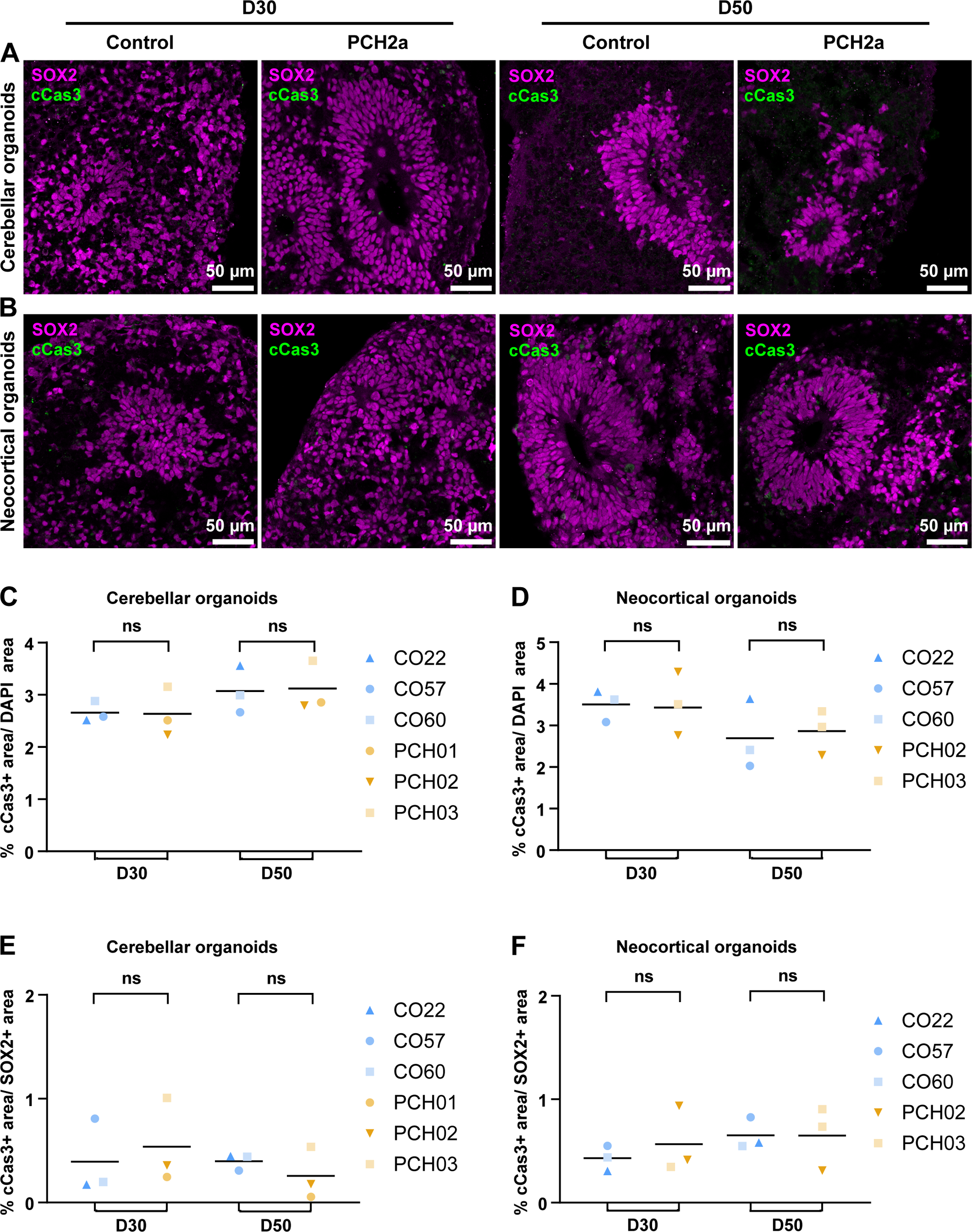
Expression of apoptotic marker cleaved Caspase 3 (cCas3) is not altered in PCH2a organoids. (A, B): Confocal microscopy images of immunohistochemistry on cerebellar (A) and neocortical (B) organoid sections at D30 and D50 of differentiation show expression of neural precursor marker SOX2 and apoptotic marker cCas3 in rosette-like structures of organoids. (C, D) Quantification of cCas3 positive area over DAPI signal shows no significant difference in cCas3 expression between PCH2a and control in cerebellar (C) and neocortical (D) organoids at D30 and D50 of differentiation. (E, F) Quantification of cCas3 positive area within SOX2+ area shows no significant difference in cCas3 expression between PCH2a and control in cerebellar (E) and neocortical (F) organoids at D30 and D50 of differentiation (p>0.05 unpaired t-test with Welch’s correction assuming unequal SDs) Representative images: Cerebellar organoids CO22 P19, PCH03 P19 (D30/D50), neocortical organoids CO57 P18, PCH02 P17 (D30/50).

### PCH2a cerebellar organoids show earlier establishment of dense SOX2+ structures neocortical organoids demonstrate no difference in SOX2+ structures

An alternative hypothesis that could explain the reduced size of cerebellar and neocortical organoids is an altered proliferation of NPCs. Immunohistochemical analysis of cerebellar organoids at D30 and D50 of differentiation showed that the number and structure of SOX2+-rich rosette structures differed between PCH2a and control cerebellar organoids (Figure 6A) with no apparent differences in neocortical organoids (Figure 6B). In D30 PCH2a cerebellar organoids, SOX2+ rosettes took on average 24% of the total organoid area, whereas in control organoids, only 2% of organoid area was taken by SOX2+cells organized in rosettes (Figure 6C). At D50 this trend was reversed: control cerebellar organoids showed a higher area of SOX2+ structures over total organoid area (12%) than PCH2a organoids (2%). Interestingly, neocortical organoids did not show differences in the area of SOX2+ rosette structures over the total organoid area on both D30 and D50 of differentiation (Figure 6D). As organoids differentiated from D30 to D50, the percentage of SOX2+ rosettes increased in both PCH2a and control neocortical organoids (Figure 6D). To determine if SOX2+ rosette structures also possessed a distinct morphology in cerebellar PCH2a organoids, we quantified the average size and thickness of the SOX2+ structures. At D30 of differentiation, PCH2a cerebellar organoids had significantly bigger SOX2+ structures; this difference was reversed at D50 (Supplementary Figure 4A). Neocortical organoids did not show significant differences in the size of SOX2+ structures at D30 or D50 of differentiation (Supplementary Figure 4B). Moreover, we measured the thickness of SOX2+ zones. SOX2+ structures were significantly thicker in PCH2a cerebellar organoids at D30 of differentiation than in controls, while control cerebellar organoids demonstrated thicker structures at D50 of differentiation than PCH2a cerebellar organoids (Supplementary Figure 4C). In contrast, in neocortical organoids, at D30 the thickness of Sox2+ structures was slightly reduced in PCH2a organoids compared to controls (Supplementary Figure 4D). At D50 there was no significant difference in thickness in neocortical organoids (Supplementary Figure 4D). To investigate if the altered SOX2+ rosette structures in cerebellar and neocortical organoids also translated to higher proliferation within these structures, we quantified the proportion of Ki-67+ cells among SOX2+ cells (Figure 6E,F). By comparing of Ki-67+/SOX2+ proportion between PCH2a and control cerebellar organoids at D30, we found higher proliferation of the SOX2+ cells (Figure 6G). Conversely, proliferation of SOX2+ cells was lower in D50 PCH2a cerebellar organoids compared to control (Figure 6G). In neocortical organoids, however, we did not find any differences in the proportion of Ki-67+/SOX2+ cells between PCH2a and control in at both D30 and D50 of differentiation (Figure 6H). Taken together, these analyses indicate aberrant proliferation properties of the NPCs in brain organoids. In line with the severity of the growth deficit (Figure 3), cerebellar organoids show dramatic changes in progenitor cell properties in PCH2a compared to control, while only subtle differences were found in neocortical organoids when comparing PCH2a and control differentiations.

**Figure 6:**
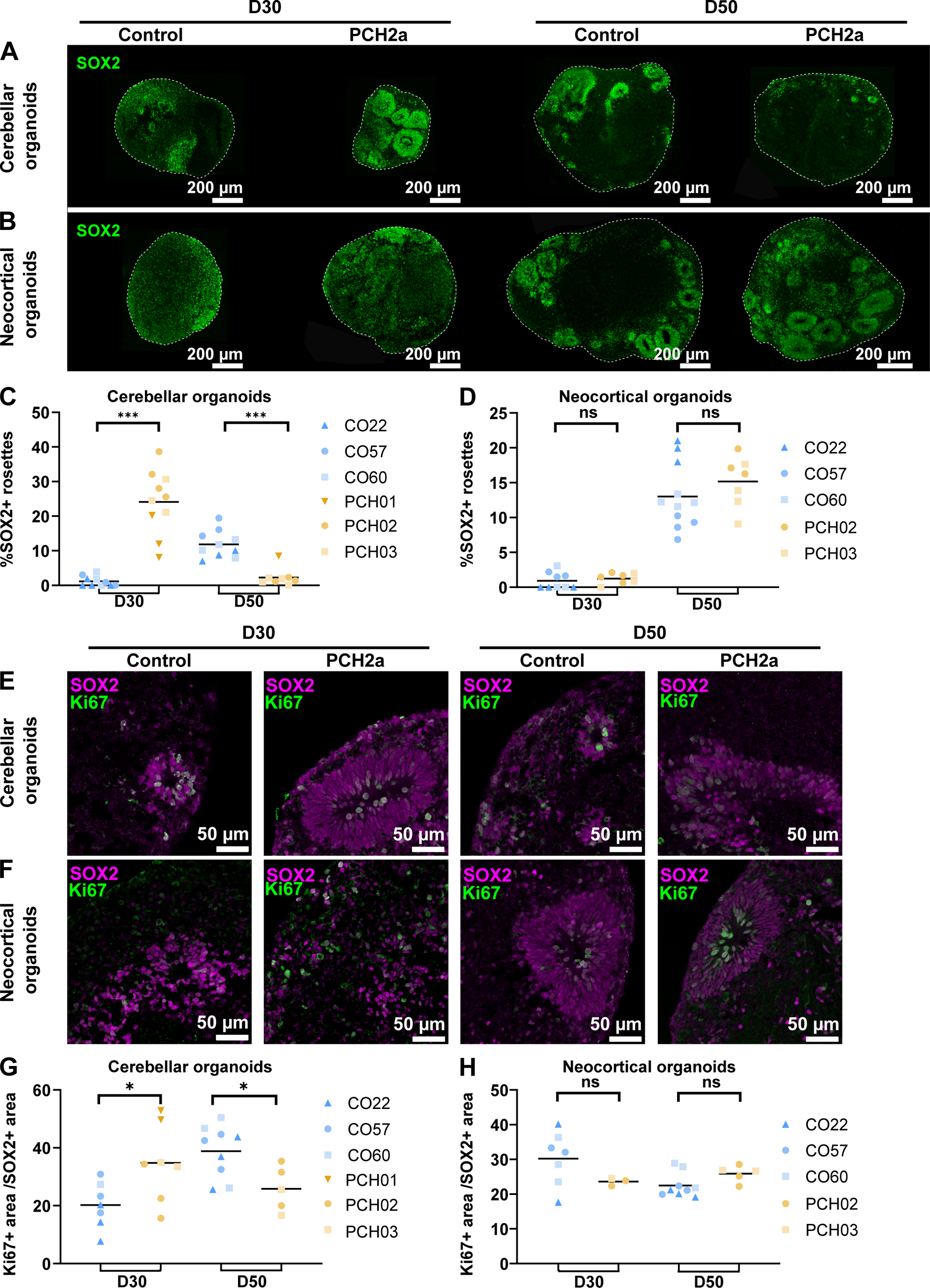
PCH2a cerebellar organoids show earlier establishment of dense SOX2+ structures while neocortical organoids demonstrate no difference in SOX2+ structures. (A+B) Epifluorescent images of immunohistochemistry on cerebellar (A) and neocortical (B) organoids at day 30 and 50 of differentiation show the expression of SOX2 (NPCs) in control (left) and PCH2a (right) organoids. Representative images: Cerebellar organoids CO57 P18, PCH02 P17 (D30), CO57 P18, PCH03 P19, neocortical organoids CO60 P19, PCH02 P17 (D30), CO22 P19, PCH02 P17 (D50). (C+D) Quantitative analysis of area covered by SOX2+ structures normalized to the area of the organoid. (C) PCH2a cerebellar organoids show significantly higher proportion of dense SOX2+ structures at D30 whereas control organoids show these structures at D50 (***, p<0.001 unpaired t-test with Welch’s correction assuming unequal SDs). (D) Neocortical organoids do not show significant differences at D30 and D50 (ns, p>0.05 in Kolmogorov-Smirnov test assuming unequal distribution). (E+F) Confocal images of immunohistochemistry against Ki-67 (magenta) and Tuj1 (green) in cerebellar (E) and neocortical (F) organoid sections at D50 illustrates the expression of Ki-67 in PCH2a (right) and control (left) organoids within SOX2+ structures. Representative images: Cerebellar organoids CO57 P18, PCH03 P19 (D30), CO57 P18, PCH03 P19, neocortical organoids CO60 P19, PCH02 P17 (D30), CO60 P19, PCH02 P17 (G+H) Quantitative analysis of percentage of Ki-67+ area normalized to SOX2+ area in cerebellar and neocortical organoids at D30 and D50 of differentiation. (G) PCH2a cerebellar organoids show significantly higher proportion of Ki-67+ area at D30 (*, p<0.05 in Kolmogorov-Smirnov test assuming unequal distribution). This difference is reversed at D50 of differentiation. (H) Neocortical organoids do not demonstrate significant differences at D30 and D50 of differentiation in Ki67+/Sox2+ area (ns, p>0.05 unpaired t-test with Welch’s correction assuming unequal SDs).

## Discussion

Understanding the cellular and molecular mechanisms of PCH2a has been hampered to date by the lack of a model replicating neuroanatomical hallmarks of the disorder such as the brain region-specific hypoplasia. Moreover, no study to date has modeled the specific variant underlying PCH2a in a neural cell. In this study, we aimed to close this gap by generating human brain region-specific organoid models of PCH2a. To achieve that, we: (1) derived three iPSC lines from affected individual (Figure 1), (2) compared these lines with control iPSC lines generated by the same protocol (Figure 1,2), and (3) extensively characterized PCH2a and control iPSC-derived cerebellar and neocortical organoids (Figures 3-6). We found that, while PCH2a-derived iPSC did not differ from control lines (Figure 2), both cerebellar and neocortical organoids demonstrated disease-relevant phenotypes. Thus, we propose that human brain region-specific organoids can serve as models to study cellular and molecular mechanisms underlying PCH2a in a tissue-specific manner.

### Using patient-derived iPSCs to model rare diseases

An important consideration in light of the current study, is the suitability of using patient-derived iPSCs for disease modeling in vitro. A major concern in this context is the reproducibility between iPSC lines derived from non-related individuals with distinct genetic backgrounds. Importantly, PCH2a is a genetically homogenous disorder, characterized by a single homozygous missense mutation and may thus be particularly suited to such an experimental design. It has been reported that the major source of variation between different iPSC lines is their genetic background (Volpato & Webber 2020). Genetic background inevitably differs between PCH2a-derived and control cell lines and thus serves as a technical confounder that may increase variation (Volpato & Webber 2020). It may also allow overlooking additional genetic variants that affect the disease course but were missed in the causal gene identification. In order to mitigate such confounding effects, we analyzed iPSC properties in great detail (Figure 2) and found no differences between PCH2a and control lines. In our organoid differentiations, we also achieve robust differences between patient and control lines despite of the genetic differences. Further, we did not find significant difference for the organoid sizes within the control or PCH2a group (Figure 3). We propose that the fact that we can generate highly reproducible data while using cell lines derived from unrelated individuals can also be seen as a strength of our study. The lack of genetic engineering using CRISPR-Cas9 also means that no off-target effects can confound our results. In conclusion, while we show consistent results between the three control lines and the three PCH2a lines, respectively, an important extension of our study, will be to confirm our findings in isogenic cell lines.

### Brain region-specific anatomical hallmarks recapitulated in organoids

In individuals affected by PCH2a, a cerebellum reduced in size is already present at birth, while a progressive reduction of cerebral volumes may be detected with time, suggesting an ongoing atrophic process of neurodegeneration (Ekert et al 2016). Analogously, we find that cerebellar PCH2a organoids are severely reduced in size from early stages on, while neocortical PCH2a organoids start displaying differences in growth at later stages of development (Figure 3). Additionally, differences in cerebellar organoid size between PCH2a and controls at later stages are larger compared to neocortical PCH2a and control organoids (Figure 3). In patients, the atrophy of supratentorial structures could be either caused by a primary effect of the disease-causing variant or as a consequence of the lack of inputs from the cerebellum, as has been discussed in very preterm infants with cerebellar lesions (Limperopoulos et al 2014). Our data indicate that the disease-causing *TSEN54* variant directly affects neocortical development. We therefore suggest that together, primary as well as secondary effects of the pathogenic variant on cerebellar and neocortical development may contribute to the diverse clinical phenotype of PCH2a. Notably, just 20% of the human cerebellum is involved in motor function (Haldipur et al 2022, Marek et al 2018), and cerebellar hypoplasia disrupting cerebellar-cerebral projections may, thus, also contribute directly to the pathological hallmarks of PCH2a not related to motor function, such as neurodevelopmental delay and lack of language development.

### Novel insights into the disease mechanism of PCH2a using the organoid models

Our model provides the foundation for studying cellular and molecular disease mechanisms underlying PCH2a as we can generate species-specific biomaterial with brain region-specific fate for subsequent analysis of the biochemical and cellular differences. Elucidating mechanisms of PCH2a has implications for PCH subtypes that are caused by variants in TSEN genes and CLP1 (Schaffer et al 2019). It can further be relevant for other rare neurological disorders caused by defects in tRNA processing machinery (Schaffer et al 2019).

Mutations in *TSEN54* lead to aberrant tRNA pools in human fibroblasts (Sekulovski et al 2021). Assuming aberrant tRNA pools are also present in the cerebellar and neocortical PCH2a organoids generated in this study, it remains elusive how altered tRNA pools translate to the reduced size in PCH2a organoids resembling the clinical phenotype of affected individuals. Interestingly, in different systems, tRNAs can directly regulate apoptosis (Avcilar-Kucukgoze & Kashina 2020, Mei et al 2010). Apoptosis has been suggested to occur in the cerebellum of affected individuals with PCH based on neuropathological observations in infants, children and adults (Barth et al 2007) as well as in animal models of PCH (Ermakova et al 2018, Kasher et al 2011, Schmidt et al 2022), respectively. We therefore investigated whether elevated apoptosis in our cerebellar and neocortical PCH2a organoids may explain the observed size differences (Figure 5). At the very early time points of differentiation in which neural progenitor cells are the dominant cell population, which are the focus of this study, we did not find any differences in apoptosis between PCH2a and control cerebellar and neocortical organoids (Figure 5). It is possible that at later stages of differentiation as neurons mature apoptosis occurs recapitulating human pathology. Future work addressing apoptosis rates in later stages of cerebellar organoid differentiation, which have recently been optimized (Chen et al 2023), will be informative to answer this question.

In our hands, cerebellar organoids show altered proliferation indicated by thickness and number of SOX2+ rosettes as well as area of SOX2+ rosettes over total organoid area, and percentage of Ki67+/SOX2+ dividing NPCs. Moreover, the direction of the changes in NPC proliferation is reversed from D30 to D50 of differentiation: PCH2a cerebellar organoids at D30 of differentiation appear to consist largely of proliferative SOX2+ cells, while they lose the majority of SOX2+ area by D50. Such a change is seemingly counterintuitive to the smaller size of the PCH2a cerebellar organoids compared to control ones over the course of differentiation. While this phenomenon may be explained through altered balance between proliferation and differentiation in the cerebellar organoids, the exact mechanism cannot be identified conclusively from our current study. Further in-depth analysis of our organoid model using, for instance, single-cell transcriptomics, will reveal cellular and molecular differences between PCH2a and control organoids to explain the differences in size.

Supporting the hypothesis that impaired proliferation and differentiation in the cerebellum leads to hypoplasia, bi-allelic variants in *PRDM13* (OMIM *6167441) cause cerebellar hypoplasia in humans (OMIM #619909) and loss of function of *prdm13* disrupts PC differentiation in zebrafish (Coolen et al 2022). Moreover, altered development in several brain structures has been reported in human brain samples of a subtype of PCH (Patel et al 2006).

Assuming tRNA pools are altered in PCH2a brain organoids as has been reported in fibroblasts (Sekulovski et al 2021), these could directly affect differentiation. Specific tRNAs can change cell state and regulate proliferation and differentiation in concert with mRNAs (Gingold et al 2014). Interestingly, in cancer, specific tRNAs can even promote metastatic progression (Goodarzi et al 2016). However, how would such changes in tRNA processing lead to brain region-specific pathology? In yeast, deficits in tRNAs are not problematic under normal conditions, however, when challenged, tRNA-depleted yeast show additional phenotypes (Bloom-Ackermann et al 2014). We, therefore, hypothesize that the developing cerebellum and, to a lesser extent, the developing neocortex have specific requirements for appropriate tRNA pools, originating perhaps from the increased neuronal output (Haldipur et al 2022, Miller et al 2019). It has been suggested that TSEN is required for processing cerebellum-specific pre-tRNAs (Sekulovski et al 2021). Indeed, tRNA isodecoders display tissue-specific expression (Ishimura et al 2014, Pinkard et al 2020) and tRNA modifications change as oligodendrocyte precursor cells differentiate into oligodendrocytes (Martin et al 2022), indicating that tRNA pools may be important regulators of neural lineage progression.

### Brain region-specific organoid models for neurogenetic disorders

Human brain organoid models have been used extensively to study neurogenetic disorders in a human cellular context (Khakipoor et al 2020, Velasco et al 2020). In most cases, cerebral organoid models have been employed, even when the cerebellum was primarily affected by the disorder (Bras et al 2022) since cerebellar differentiation protocols have been developed only recently (Hua et al 2022, Muguruma et al 2015, Nayler et al 2021, Silva et al 2020). Therefore, to date, the use of cerebellar organoids in neurogenetic disease modeling has not been demonstrated. For the first time in this study, we show how cerebellar organoids can be employed to model the brain region-specific neuropathology of PCH2a, a severe neurological disorder that primarily affects the cerebellum and pons and, to a lesser extent, the neocortex. PCH2a iPSCs robustly differentiate into both neocortical and cerebellar fate (Figures 4, 5) in line with the observation that regional specification was not affects in a *tsen54* loss-of-function zebrafish (Kasher et al 2011) and different cerebellar cell types including PCs and GCs were found in pathological studies of PCH2a (Rudnik-Schöneborn et al 2014). However, we observe brain region-specific differences in organoid growth and proliferation between PCH2a and control organoids. We therefore suggest that using organoid protocols that replicate regional identity of the affected brain region is crucial to study a specific neuropathology. Consequently, if a disease affects multiple brain regions, the full potential of organoid technology may only be realized by combining different brain region-specific organoids.

## Acknowledgements

We thank the affected individuals and their families for donating samples. We would like to thank PCH-Familie e.V. for their support, especially Julia Matilainen and Axel Lankenau, for the fruitful discussions. We thank Clemens Lumper, Elisabeth Gustafsson, Lea Fischer, Jasmin Treu, Maximilian Feige, Christina Kulka, Ezgi Atay, Felix Hildebrand, Melanie Kraft, and Yvonne Schelling for technical support. We thank Nicolas Snaidero for support with confocal microscopy. We thank Javier Martinez, Stefan Weitzer, and Hansjürgen Volkmer for critical feedback on the manuscript.

## Contributions

TK designed the study, performed experiments, data analysis, and statistical analysis, and prepared the manuscript; SH generated iPSC lines; KS performed experiments, statistical analysis and prepared the manuscript; KB performed experiments and data analysis; ZY performed experiments; LL, SG, WJ, and IKM provided clinical expertise and data; LS supervised the generation of iPSC lines; all authors revised the manuscript; SM conceived and designed the study, supervised the work, and prepared the manuscript.

## Methods

### Recruitment of affected individuals

Affected individuals were recruited within our PCH2 natural history study, collecting clinical and diagnostic data, including diagnostic MR images. Written informed consent was obtained from guardians and archived. All procedures were performed in accordance with the Helsinki Declaration. Individual-level data were de-identified. The study was approved by the ethics committee of the medical faculty, the local Institutional Review Boards of the Medical Faculty of the University of Tübingen, Germany (961/2020BO2 and 598/2011BO1) and Freiburg, Germany (20-1040).

### Diagnostic confirmation by genetic sequencing

Next-generation sequencing and/or Sanger sequencing was performed after obtaining written informed consent for either clinical sequencing and/or center-specific institutional review board-approved research sequencing. All affected individuals harbored the hypomorphic founder variant c.919G>T in *TSEN54* in the homozygous state. The bi-allelic localization was confirmed by carrier testing.

### Skin biopsies

Skin biopsies were acquired at different ages (9 months to 15 years) according to local standards of routine diagnostic procedures.

### Culturing and reprogramming fibroblasts

Human dermal fibroblasts were obtained from skin biopsies and cultivated in Dulbecco’s modified eagle medium (DMEM) (Thermo Fisher Scientific) supplemented with 10% fetal bovine serum (FBS) (Thermo Fisher Scientific) (fibroblast medium).

iPSC generation from fibroblasts was performed according to a published protocol with minor modifications (Okita et al 2013). Briefly, reprogramming was initiated by nucleofection of 1×10^5^ fibroblast with 1 µg of each episomal plasmid (pCXLE-hUL, pCXLE-hSK and pCXLE-hOCT4) using the Nucleofector 2b (Lonza). Initially, fibroblasts were cultivated in fibroblast medium supplemented with 2 ng/ml FGF2 (Peprotech, Cat. no. 100-18B). On D3, the medium was changed to Essential 8 (E8) medium containing 100 µM sodium butyrate (NaB, Sigma-Aldrich, Cat. no. B5887). After 3 – 4 weeks, with media changes every other day, iPSC colonies were manually picked and further expanded in E8 medium performing media changes daily. After ≥5 passages, they were genomically and functionally characterized and frozen in E8 medium containing 40% KO-SR (Thermo Fisher Scientific, Cat. no. 10828-028), 10% DMSO (Sigma-Aldrich, Cat. no. D4540) and 1 µM Y-27632 (Selleck Chemicals, Cat. no. S1049). All iPSC lines used in this study (3 control lines, 3 PCH2a lines) were characterized according to the scientific guidelines for Lab Resources (Stem Cell Research).

**Table 1:** Origin of iPSC lines.

| Cell line ID | Donor age | Ethnicity | Sex | Pathological Variant |
| --- | --- | --- | --- | --- |
| CO22 | 74 years | European | male | N/A |
| CO57 | 46 years | European | male | N/A |
| CO60 | 22 years | European | male | N/A |
| PCH01 | 9 months | European | male | c.919G>T |
| PCH02 | 14 years | European | male | c.919G>T |
| PCH03 | 4 years | European | male | c.919G>T |

### Genomic integrity analysis

In order to verify genomic integrity, DNA of iPSCs and fibroblasts was isolated with DNeasy Blood & Tissue Kit (Qiagen) according to the manufacturer’s guidelines. Whole-genome SNP genotyping (SNP array) was conducted using Infinium OmniExpressExome-8-BeadChip (Illumina) and GenomeStudio V2.0.3 (Illumina) for evaluation. Copy number analysis was performed using CNVPartition plugin (Illumina). Early mosaicism states were evaluated by manual review of B allele frequency plots on a chromosomal level. Results can be found in the supplementary data.

### Pluripotency assessment

To assess the alkaline phosphatase (ALP) expression or for immunocytochemical analysis, iPSCs were fixed with 4% paraformaldehyde (PFA) and either assessed for ALP expression or permeabilized with 0.1% Triton X-100, blocked with 5% FBS and stained overnight at 4°C with primary antibodies for immunocytochemical analysis (rabbit anti-OCT4, 1:100, Proteintech, Cat. no. 11263-1-AP / mouse anti-TRA-1-81, 1:500, Millipore, Cat. no. MAB4381). Samples were visualized after staining with Alexa Fluor 488-conjugated secondary antibodies (Thermo Fisher Scientific) for 1h at room temperature. Nuclei were counterstained with Hoechst 33342 (1:10.000, Invitrogen). Samples were embedded in ProLong Gold Antifade Reagent (Thermo Fisher Scientific, Cat. No. P36930) and imaged with AxioImager Z1 (Zeiss).

The differentiation capacity of iPSCs into cells of all three germ layers was determined by an embryonic body (EB)-based protocol. 1.2×10^6^ iPSCs were seeded in AggreWell800 plates (Stem cell technologies) in EB medium consisting of DMEM/F-12 supplemented with 20% Knockout Serum Replacement, 1% MEM Non-essential-amino-acid solution, 1% Pen/Strep, 1% GlutaMAX and 50µM β-Mercaptoethanol. On D4, EBs were plated onto coverslips for further differentiation. Specific marker expression (TUJ (mouse anti-TUJ, 1:1,000, Sigma Aldrich, Cat. no. T8660), SMA (mouse anti-SMA, 1:100, Dako, Cat. no. M0851) was assessed after 10 days as described above. For endodermal induction of iPSCs, 2×10^5^ cells were seeded onto coverslips and cultivated in endoderm induction medium consisting of RPMI1640 advanced supplemented with 1xB27, 1% Pen/Strep, 0.2% FCS, 2µM CHIR-99021 and 50ng/ml Activin A. At D4 of differentiation, cells were stained for FOXA2 (rabbit anti-FOXA2, 1:300, Millipore, Cat. no. 07-633), following the fixation and immunocytochemistry protocol mentioned above.

### Sanger sequencing

To ensure the correct genetic background of all iPSC lines and control, Sanger sequencing was performed. Genomic DNA was extracted using DNA isolate kit (BioCat, Cat. No. BIO-52066-BL). The region of interest was amplified using Phusion® High-Fidelity PCR Kit (New England Biolabs, Cat. No. E0553S). The PCR product was purified with QIAquick PCR Purification Kit (Qiagen, Cat. No. 28104), and samples were sent to Eurofins for sequencing. The following primers were used: Forward (AGAAACCCCAGGAGT), reverse (CTCAATCCATCCGAG).

### iPSC culture

iPSC lines derived from affected individuals and control lines were generated following the same protocol and cultured under standard conditions (37°C, 5% CO2, and 100% humidity) in E8 Flex medium (Gibco, Cat. no. A2858501) on hESC-qualified growth factor-reduced Matrigel-coated (Corning, Cat. no. 354277) cell culture dishes (Greiner, Cat. no. 657160). Passaging was performed in colonies using Gentle Dissociation Reagent (STEMCELL Technologies, Cat. no. 07174) once the culture reached 80%-90% confluency. The culture medium was supplemented with Thiazovivin (Sigma-Aldrich, Cat. no. 420220) until the following day. All cell lines were tested for mycoplasma contamination regularly with PCR Mycoplasma Detection Set (TaKaRa, Cat. no. 6601) and maintained under passage 20. The pluripotency for each cell line was confirmed with antibodies against OCT4 (rabbit, 1:500, Abcam, Cat. no. ab19857) before each differentiation.

### Staining of iPSCs

In order to assess the pluripotency and proliferation of all iPSCs used in this study 3 consecutive passages per cell line were cultured on coverslips, fixed and stained for OCT4 (rabbit, 1:500, Abcam, Cat. no. ab19857), Ki67 (rabbit, 1:400, CST, Cat. no 9661S) and cCas3 (rabbit, 1:600, Merck, Cat. no AB9260). The cells were fixed with 4% paraformaldehyde (PFA, Morphisto, Cat. no. 11762) in PBS for 15 minutes and carefully washed twice with 1x PBS (Roth, Cat. no. 1105.1). Prior to staining, the cells were permeabilized with 0.5% Triton X-100 (Sigma, Cat. No. T8787) for 10 minutes at room temperature. After washing the cells with 1x PBS for 5 minutes, they were incubated in blocking buffer, consisting of 10% Normal Donkey Serum (NDS, Abcam, ab7475) in PBS, for 1 hour at room temperature. The primary antibodies were diluted in blocking buffer and administered to the cover slips over night at 4°C. The cells were washed three times with 1x PBS for 5 minutes and incubated with the secondary antibodies in blocking buffer for 1 hour at room temperature. After two washes with 1x PBS for 5 minutes, the cells were counterstained with DAPI (1:5000) (Thermofisher Scientific, Cat. no. D1306) in PBS. Finally, the cells were washed once in 1x PBS and mounted on slides using ProLong Gold (Thermofisher Scientific, Cat. no. P36930). For all cell lines and conditions coverslips were imaged at 20x magnification.

### EdU incorporation

To quantify iPSC proliferation 3 passages per cell line were treated with 10 µM EdU (Thermo Fisher Cat. no. C10338) 3-4 days after last passage for 1 and 4 hours. After the incubation with EdU cells were fixed and click-chemistry was performed as advised by the manufacturer. EdU signal was labeled with provided Alexa 555 dye. Nuclear staining was performed with DAPI (Thermofisher Scientific, Cat. no. D1306). Coverslips were imaged at 20x magnification on a confocal microscope, keeping laser settings identical for all cell lines and conditions.

### Quantification of immunohistochemistry and click-chemistry on iPSCs

In order to quantify possible differences in the expression of markers for pluripotency, proliferation, apoptosis and the number of cells positive for EdU we stained and imaged respective samples in one experiment. All samples were imaged with the same laser intensity settings on a confocal microscope. Raw image files were further processed in FIJI (Schindelin et al 2012). Here, we used the watershed algorithm on the DAPI channel to identify individual nuclei. These were then registered as regions of interest (ROIs) and counted. To analyze the number of stained cells, thresholding the respective channel of interest was performed with identical parameters for each channel and all samples according to the negative staining control. ROIs demonstrating a signal for the channel of interest were then measured and counted. This allowed us to analyze the percentage of cells positive for the marker of interest within the total population of cells. Finally, statistical analysis and plotting were performed in GraphPad Prism.

### Generation of cerebellar organoids

Cerebellar organoids were generated as previously described (Silva et al 2020) with some alterations: 80-90% confluent iPSCs were dissociated into single cells using Accutase (Merck, Cat. no. A6964), and 4,500 cells were seeded per well of 96 well V-bottom low adhesion plates (S-bio, Cat. no. MS-9096VZ) in E8 Flex medium (Gibco, Cat. no. A2858501), supplemented with 10 μM Y-27632 (Cayman Chemical, Cat. no. 10005583). Once the aggregates reached a diameter of 250 μm, the medium was changed to growth factor-free chemically defined medium (gfCDM), supplemented with 50 ng/ml FGF2 (PeproTech, Cat. no. 100-18B) and 10 μM SB-431542 (Tocris, Cat. No. 1614). At D7 of differentiation, FGF2 and SB-431542 were reduced to 33.3 ng/ml and 6.67 μM, respectively. At D14, media was supplemented with 100 ng/ml FGF19 (PeproTech, Cat. No. 100-32). The medium was changed to Neurobasal Medium at D21, supplemented with 300 ng/ml SDF-1 from D28 to D34. From D35 onwards, media was changed to complete BrainPhys (StemCell Technologies, Cat. No. 5793), supplemented with 10 μg/ml BDNF (PeproTech, Cat.No. 450-02), 100 μg/ml GDNF (PeproTech, Cat.No. 450-10), 100 mg/ml dbcAMP (PeproTech, Cat.No. 1698950) and 250 mM ascorbic acid (Tocris, Cat. No. 4055).

### Generation of cortical spheroids

Cortical spheroids were generated as previously described (Pasca et al 2015) with only minor alterations. In brief, 80-90% confluent iPSCs were dissociated into single cells using Accutase (Merck, Cat. no A6964) and 9000 cells were seeded per well of 96 well V-bottom low adhesion plates (S-bio, Cat. no. MS-9096VZ) in E8 Flex medium (Gibco, Cat. no. A2858501) supplemented with 10 μM Y-27632 (Cayman Chemical, Cat. no. 10005583). The medium was changed to neural induction medium (NI) (Essential 6 Thermo Fisher Cat. No. A151640), supplemented with 2.5 μM Dorsomorphin (Tocris, Cat. No. 3093), 10 μM SB-431542 (Tocris, Cat. No. 1614) and 2.5 μM XAV-939 (Tocris, Cat. No. 3748) the next day. NI medium was changed every other day and replaced with neural maintenance medium (NM) (Neurobasal-A (Gibco, Cat. no.10888-022), B27-VitA (Thermo Fisher, Cat. no. 12587010), GlutaMAX (Thermo Fisher, Cat. no. 35050038), Pen/Strep (Sigma, Cat. no. P0781) at D6. NM was supplemented with 20 ng/ml EGF (Merck, Cat. No GF144) and FGF2 (PeproTech, Cat. No. 100-18B) from D6 to D24 and with 20 ng/ml BDNF (PeproTech, Cat. No. 450-02) and NT-3 (PeproTech, Cat. No. 450-03) from D25 to D43. NM was changed every other day and not supplemented after D43.

### Size measurements

To investigate the size of organoids, brightfield images of the organoids were taken at D0, D10, D20, D30, D50, D70 and D90 of differentiation with an EVOS cell imaging system (Thermo Fisher). These images were analyzed using a published macro (Ivanov et al 2014) for FIJI (Schindelin et al 2012). The data were further analyzed with Excel, and Graph Pad Prism was used to plot data.

### Fixation, cryosections and immunohistochemistry

Organoids were fixed at respective time points in 4% paraformaldehyde (PFA, Morphisto, Cat. no. 11762) in PBS for 45–60 min at room temperature (Lancaster & Knoblich 2014). The organoids were washed three times for 15 minutes with 1x PBS (Roth, Cat. no. 1105.1) and then incubated in 30% sucrose (Sigma Aldrich, Cat. no. S7903) in PBS solution at 4°C until they sunk to the bottom of the dish. The organoids were embedded in a 1:1 v/v mixture of 30% sucrose in PBS and optimal cutting temperature (OCT) compound (Sakura, Cat. no. 4583) and sectioned on Superfrost Plus slides (R. Langenbrinck GmbH, Cat. no 03-0060) with a cryostat at 20 µm (Leica). The slides were stored at −80°C.

For immunohistochemistry, slides were thawed for 15 minutes at room temperature and the embedding solution was rinsed off with PBS. Antigen retrieval was achieved by immersing the slides in 10 mM citric acid buffer (pH 6.0) and boiling for 20 minutes in a microwave. A hydrophobic pen (PAP pen, Abcam, Cat. no. ab2601) was used to circle the sections to prevent the blocking solution from spilling during incubation. Permeabilization and blocking were performed with 1% Triton-X100 (Sigma, Cat. No. T8787), 0.2% gelatin (Sigma, Cat. no. G1890), and 10% normal donkey serum (Abcam, Cat. no. ab7475) in PBS for 1 hour at room temperature. Primary antibodies were diluted in permeabilization and blocking solution and applied to the sections overnight at 4°C. Subsequently, the slides were rinsed with PBS three times for 15 minutes, then secondary antibodies were diluted in permeabilization and blocking solution and applied for three hours at room temperature. The sections were rinsed in PBS three times for 15 minutes, and nuclei were stained with DAPI (1:5000) (Thermofisher Scientific, Cat. no. D1306) diluted in PBS for 4 minutes, rinsed in PBS and mounted using ProLong Gold (Thermofisher Scientific, Cat. no. P36930).

**Table 2:** Primary Antibodies.

| Antibody | species | vendor | cat number | dilution |
| --- | --- | --- | --- | --- |
| ATOH1 | mouse | Merck | WH0000474M1 | 1:500 |
| BAHRL1 | rabbit | Atlas Antibodies | HPA004809 | 1:500 |
| Calbindin | mouse | Merck | C9848 | 1:500 |
| cCas3 | rabbit | CST | 9661S | 1:400 |
| CTIP2 | rat | Abcam | ab18465 | 1:500 |
| Ki67 | rabbit | Merck | AB9260 | 1:600 |
| Kirrel2 | rabbit | Atlas Antibodies | HPA071587 | 1:500 |
| LHX2 | goat | Merck | SAB2500593 | 1:500 |
| Map2 | rabbit | Abcam | ab32454 | 1:500 |
| Oct4 | rabbit | Abcam | ab19857 | 1:500 |
| Olig2 | rabbit | Merck | AB9610 | 1:500 |
| SATB2 | rabbit | Abcam | ab34735 | 1:500 |
| SKOR2 | rabbit | Atlas Antibodies | HPA046206 | 1:500 |
| SOX2 | goat | R&D Systems | AF2018 | 1:500 |
| TSEN54 | rabbit | Invitrogen | PA5-101939 | 1:500 |
| Tuj1 | mouse | Atlas Antibodies | AMAb91394 | 1:500 |

**Table 3:** Secondary antibodies.

| Host species | Target species | Em wavelength | Provider | Cat.no |
| --- | --- | --- | --- | --- |
| donkey | goat | AF555 | Abcam | ab150130 |
| donkey | goat | AF647 | Abcam | A21447 |
| donkey | mouse | AF568 | Abcam | ab175472 |
| donkey | mouse | AF647 | Abcam | A31571 |
| donkey | rabbit | AF488 | Abcam | A21206 |
| donkey | rabbit | AF546 | Abcam | A10040 |
| donkey | rabbit | AF647 | Abcam | A31573 |
| donkey | sheep | AF488 | Abcam | A11015 |
| donkey | sheep | AF647 | Abcam | A21448 |

### Quantification of SOX2+ zones

To quantify SOX2-positive area within individual organoids, we stained and imaged respective samples in one experiment. The laser intensity of the confocal microscope was adjusted according to the negative staining control. All samples were imaged with the same laser intensity settings. Raw image files were further processed in FIJI (Schindelin et al 2012). To analyze the area of SOX2+ zones, these regions were manually selected and measured. For each organoid and timepoint all SOX2+ zones were quantified using the “measure” tool in FIJI. The area of the SOX2+ zones was then normalized to the total DAPI+ area of the organoid. To measure the thickness of the SOX2+ zones the outer and inner circumference was measured and the difference was calculated by subtracting the radius of the inner circumference from the radius of the outer circumference. Finally, statistical analysis and plotting were performed in GraphPad Prism.

### Quantification of cCas3 over SOX2 staining

To quantify cCas3 positive area within regions of interest in individual organoids, we stained and imaged respective samples in one experiment. The laser intensity of the confocal microscope was adjusted according to the negative staining control. All samples were imaged with the same laser intensity settings. Raw image files were further processed in FIJI (Schindelin et al 2012). To analyze the area of stained cells, thresholding for cCas3 and SOX2 channels was performed with identical parameters for each channel and all samples. First, the area of cCas3 positively stained regions was quantified using the “measure” tool in FIJI. Then the cCas3 area was normalized to the SOX2-positive area of the respective image. Finally, statistical analysis and plotting were performed in GraphPad Prism.

### Quantification of Ki67+ cells over SOX2+ cells

To quantify Ki67+/SOX2 cells within regions of interest (ROIs) in individual organoids, we stained and imaged respective samples in one experiment. The laser intensity of the confocal microscope was adjusted according to the negative staining control. All samples were imaged with the same laser intensity settings. Raw image files were further processed in FIJI (Schindelin et al 2012). Thresholding was adjusted to respective channel and for each channel identical parameters were used for all samples. Watershed algorithm was used on thresholded SOX2 channel to identify ROIs. ROIs were saved in ROI manager and results and transferred to thresholded Ki67 channel. Number of Ki67+ cells within these ROIs were counted and saved in the results file (Supplementary Figure 5). Results were exported to Excel to calculate percentage and statistical analysis and plotting were performed in GraphPad Prism.

### Statistical analysis

Details of specific statistical comparisons are listed in the relevant figure legends. No formal comparison of variances was performed between experimental groups. No statistical methods were used to predetermine sample size.

---

**Supplementary Figure 1: Size differences of cerebellar and neocortical organoids at different stages of differentiation.** Quantification of the area of cerebellar (A-C) and neocortical (D-F) control and PCH2a organoid areas at D30, D50 and D90 of differentiation. Sizes of organoids differ significantly between PCH2a and control organoids. (****, p< 0.0001, *** p<0.001 unpaired t-test with Welch’s correction assuming unequal SDs). n>8 organoids per cell line, time point and differentiation. (G) Linear regression model for growth curves of cerebellar organoids. (H) Linear regression model for growth curves of neocortical organoids.

**Supplementary Figure 2: PCH2a and control cerebellar organoids show differentiation into cerebellar lineage.** (A-D) ATOH1 (green) expression demonstrates the presence of rhombic lip derived granule cell precursors while SKOR2 (magenta) is marker for young Purkinje cells derived from the cerebellar VZ, both markers are present in control (A, B) and PCH2a (C, D) cerebellar organoids (representative images CO57 P18, PCH03 P19). Dotted lines indicate the outline of organoids, white squares (A,C) indicate the region of the zoom-in (B, D). (E, F) Staining for early Purkinje marker OLIG2 (green) and LHX2 (magenta), early marker for nuclear transitory zone cells in Day 30 control (E) and PCH2a (F) cerebellar organoids (representative images CO60 P19, PCH01 P19). Nuclei are stained with DAPI.

**Supplementary Figure 3: PCH2a and control neocortical organoids express layer-specific neuronal markers.** (A-F) Epifluorescent microscopy images of immunohistochemistry on neocortical organoid sections. Control (A) and PCH2a (B) neocortical organoids express early-born deep-layer neuronal marker CTIP2 (magenta) at D50 of differentiation (representative images CO22 P19, PCH03 P19). (C, D, E, F) Control (C) and PCH2a (D) neocortical organoids show expression of CTIP2 as well upper layer neuronal marker SATB2 (green) at D70 (C, D) (representative images CO22 P19, PCH02 P17) and D90 (E, F) (representative images CO57 P18, PCH03 P19) of differentiation (CTIP2 green, SATB2 magenta). Nuclei are stained with DAPI.

**Supplementary Figure 4: Morphology of SOX2+ structures differs in PCH2a and control organoids.** (A) SOX2+ structures are significantly bigger in PCH2a organoids at D30 of differentiation. At D50 control organoids show significantly bigger SOX2+ structures. (B) SOX2+ structures do not show difference in size at D30 and D50 of differentiation in neocortical organoids. (C) PCH2a cerebellar organoids show thicker SOX2+ structures at D30 of differentiation while SOX2+ structures are thicker in control organoids at D50. (D) Thickness of SOX2+ structures is lower in PCH2a neocortical organoids at D30 of differentiation. There is no difference in thickness of these structures at D50 of differentiation in neocortical organoids (p>0.05 for ns and p<0.05 for *, p<0.01 for**, p<0.001 for ***unpaired t-test with Welch’s correction assuming unequal SDs).

**Supplementary Figure 5: Overview of methods for image analysis for SOX2+ rosettes.** (A) Overlay of input image for size and thickness analysis of SOX2+ rosettes. (B) SOX2 channel of input image. White dotted line indicates outline of rosette used for size analysis of rosette structures. (C) SOX2 channel of input image. Outer dotted line indicates outline of SOX2+ rosette, while inner dotted structures indicates boarder to lumen. Thickness (d) of SOX2+ rosettes was calculated by subtracting the inner border from the circumference resulting in the mean thickness of the whole rosette structure. (D) Individual channels and overlay of input image for quantification of Ki67+/SOX2+ cells. (E) White dotted lines indicate region of interest (ROI) for quantification of Ki67+/SOX2+ cells. (F) Input SOX2 channel. (G) Watershed algorithm on thresholded SOX2 image. (H) Overlay of ROIs resulting from watershed algorithm on input image. (I) Ki67 input image. (J) Thresholded Ki67 image. (K) Overlay of SOX2 ROIs on thresholded Ki67 image. All SOX2 ROIs with Ki67 signal were counted as Ki67+/SOX2+ cells. Nuclei are stained with DAPI.

