## Supplementary figures and images for "Human organoid model of PCH2a recapitulates brain region-specific pathology"

### Supplementary figure 1

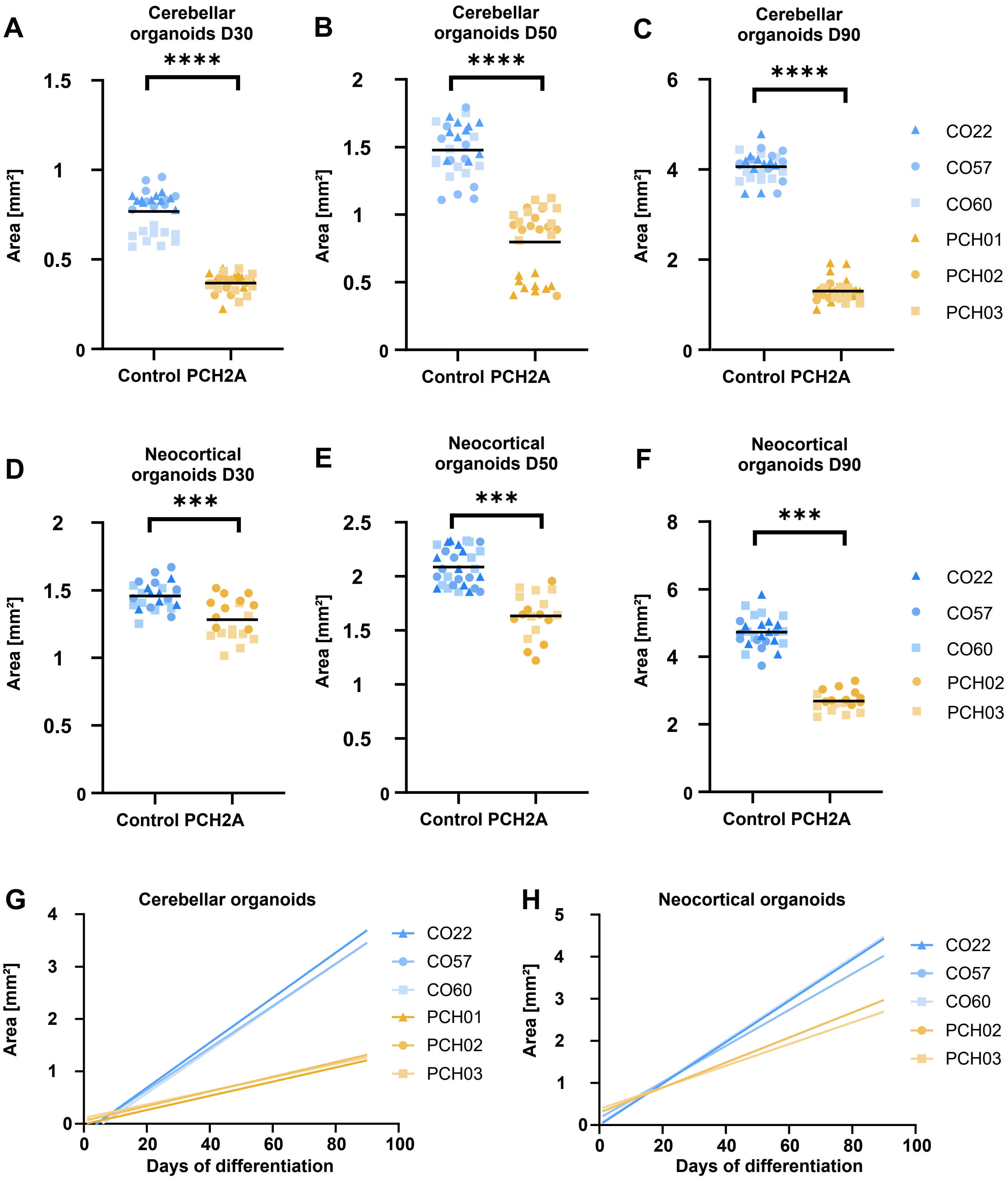

### Supplementary figure 2

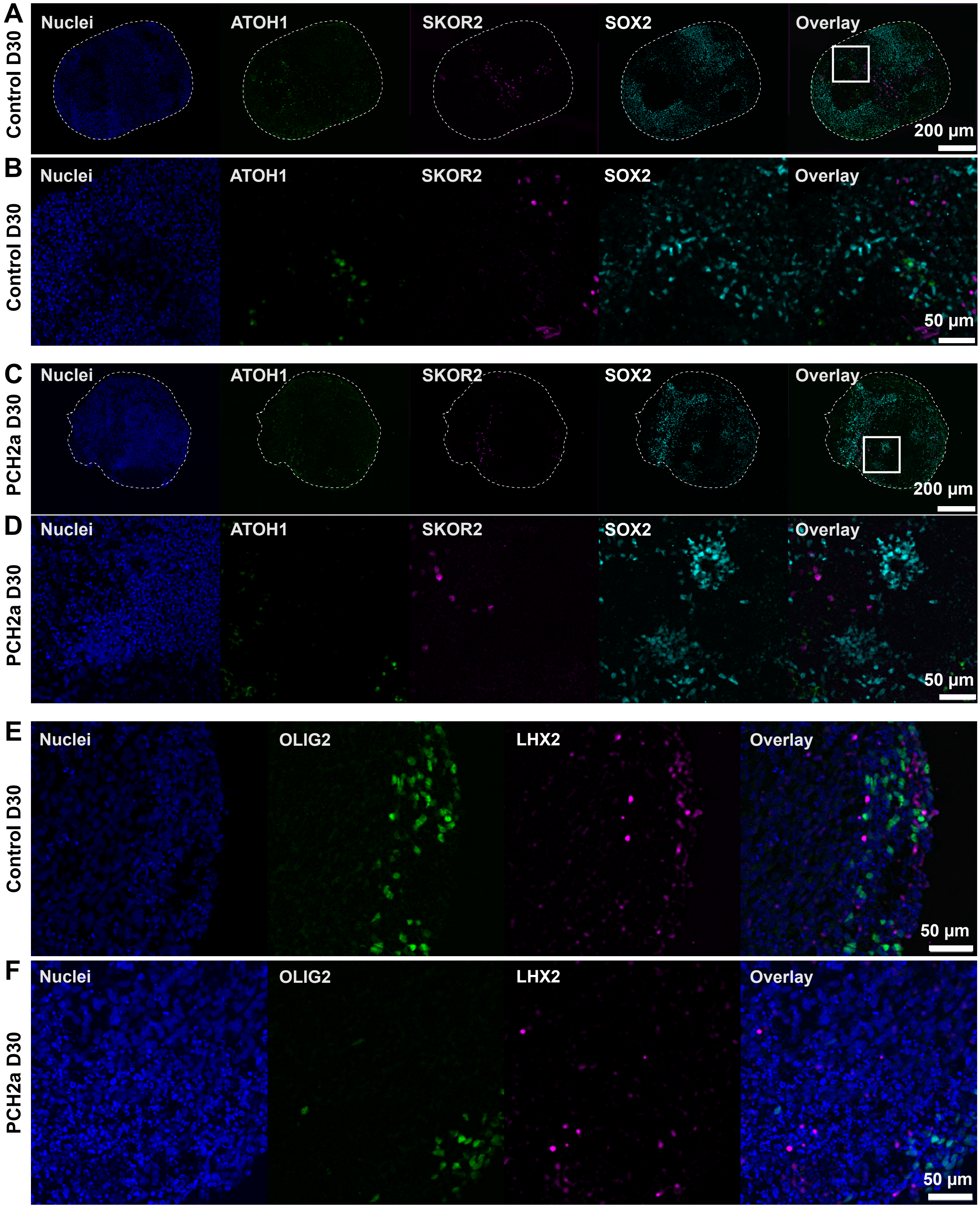

### Supplementary figure 3

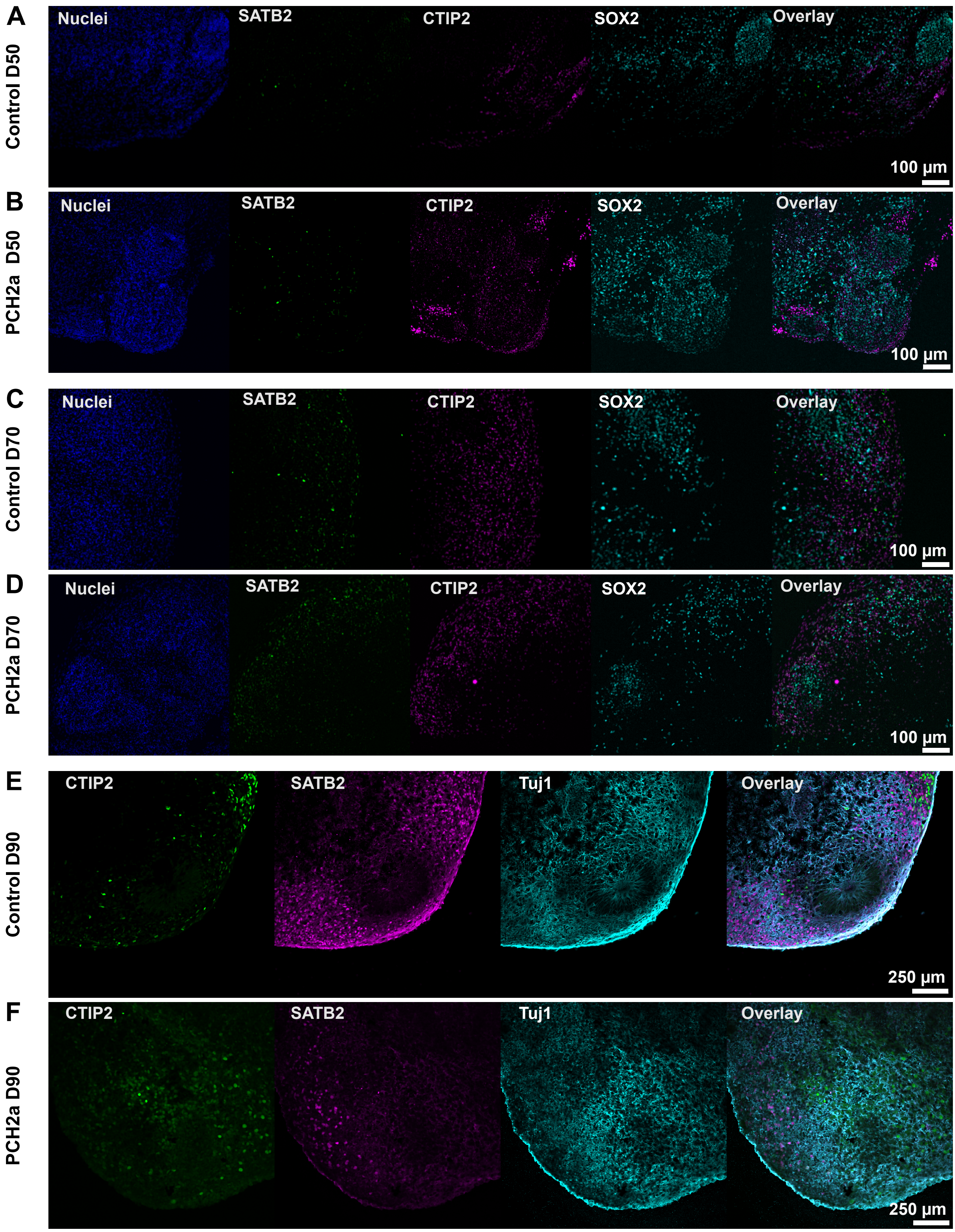

### Supplementary figure 4

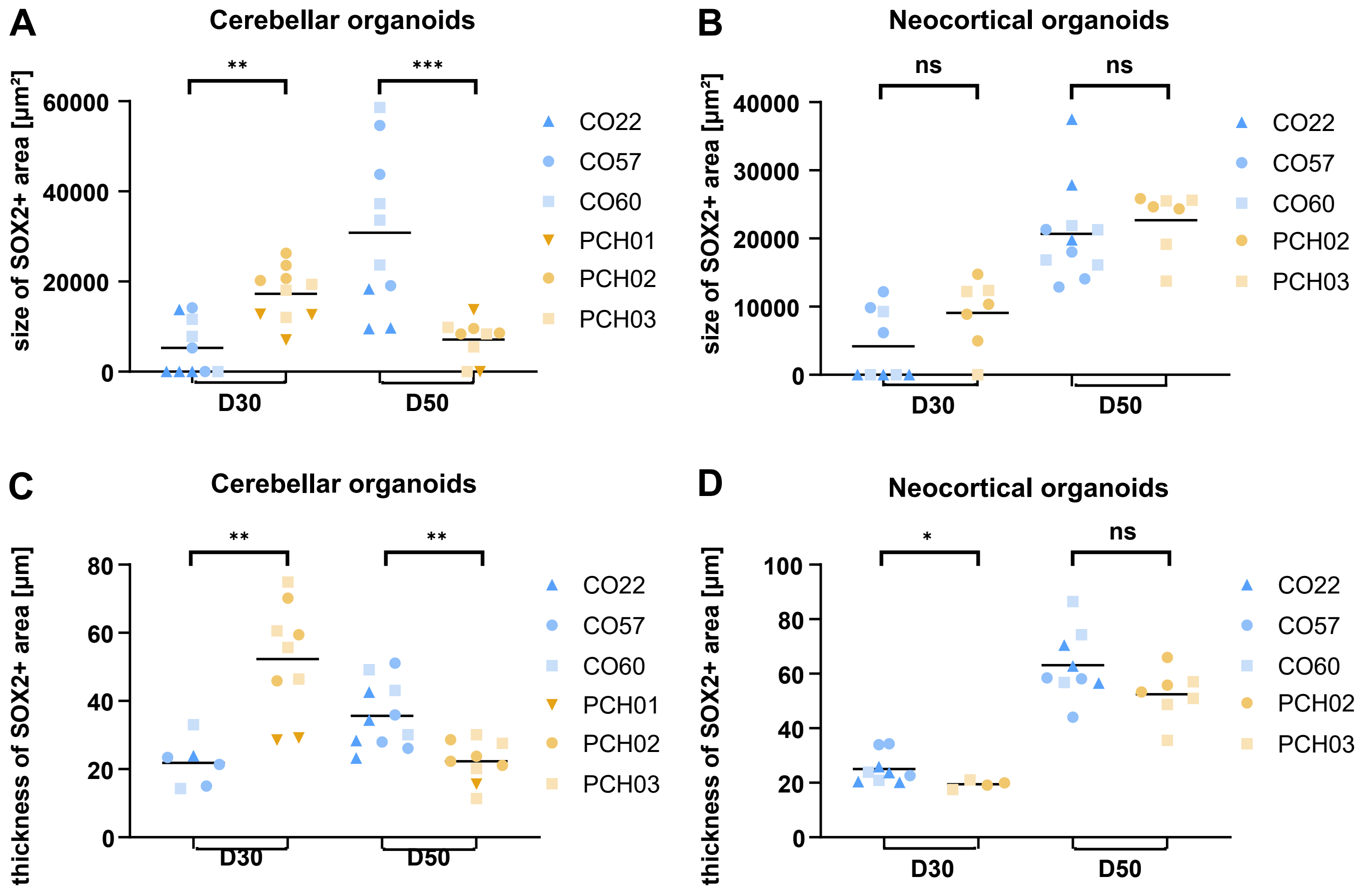

### Supplementary figure 5

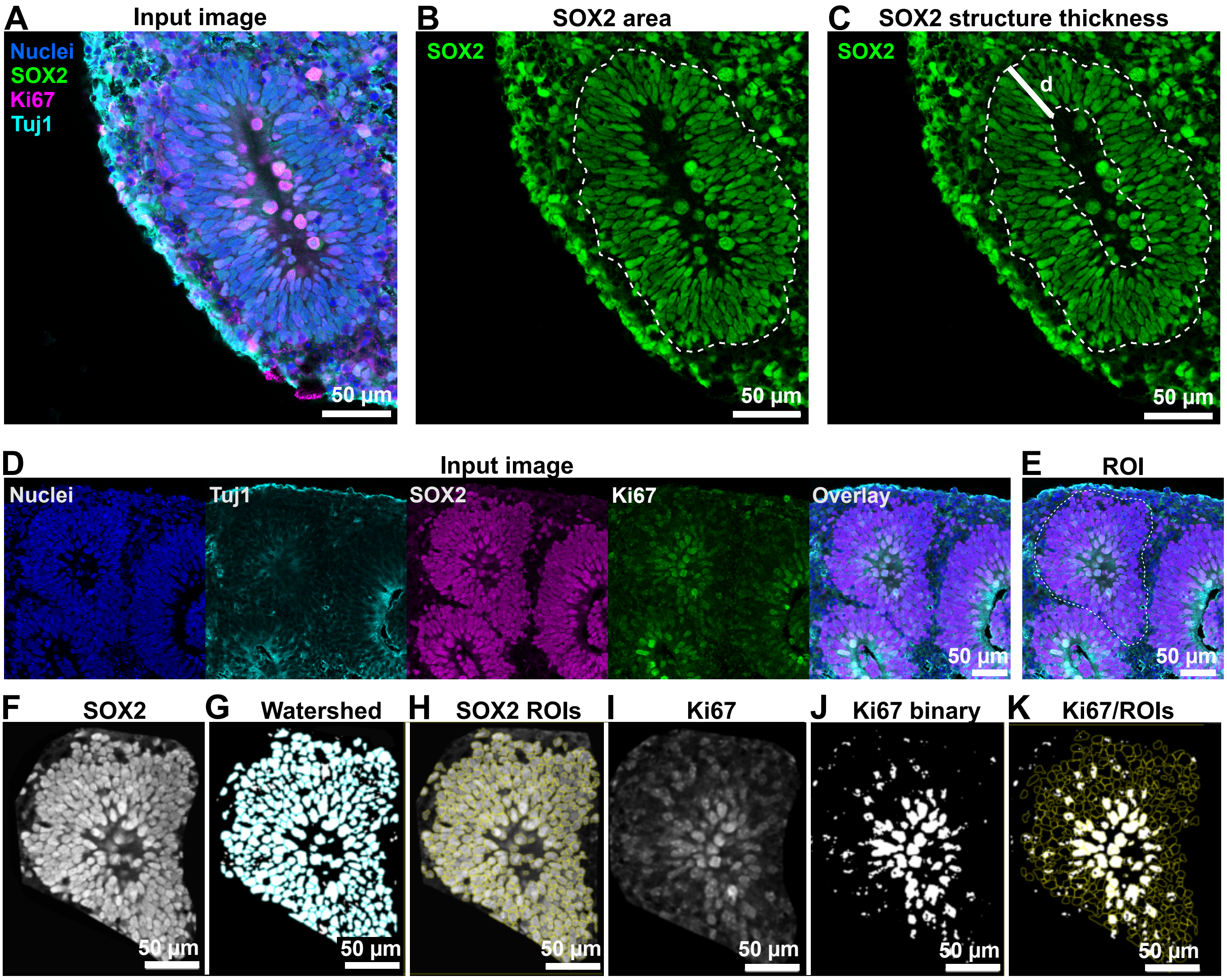
